# Temporal models using environmental variables to predict *Aedes aegypti* oviposition activity in a temperate region of Argentina

**DOI:** 10.1101/816421

**Authors:** Elisabet M. Benitez, Elizabet L. Estallo, Marta Grech, Maria Frías-Céspedes, Walter R. Almirón, Francisco F. Ludueña-Almeida

## Abstract

Environmental variables are some of the factors that more impact on *Aedes aegypti*, vector of Dengue, Chikungunya and Zika viruses. In this study, the *Ae. aegypti* oviposition activity was related to satellite and meteorological variables, in Córdoba City (Argentina). Eggs were collected using ovitraps placed throughout the city from 2009 to 2012, replaced them weekly. Generalized Linear Mixed Models were developed with negative binomial distributions of errors, modeling average number of eggs collected weekly as a function of satellite and meteorological variables with time lags. The best model included a vegetation index, vapor pressure of water, precipitation and photoperiod, lagged between 3 and 4 weeks. By each increment unit in vegetation index, vapor pressure (hPa) and light hour, the average number of eggs increases by 72%, 20% and 29%, respectively, in the following 3 or 4 weeks. In addition, the average number of eggs decreases a 50% in the following month by each millimeter of rain. The results evince the relevant importance of the vegetation, maybe due to the shade that provide to the containers as breeding habitat of this species and the shelter for adults. On the other hand, the negative effect of precipitation could be a consequence of abundant rainfalls that fulfill containers avoiding females to lay eggs in there. Although no significant effect of the photoperiod on the vector abundance has been detected in the north of the country, in Córdoba City an important effect is observed when it is presented together with other variables, producing different effects of these variables in different regions. Lastly, it is necessary to emphasize the importance of the minimum temperature in temperate zones, since it would be a limiting factor that prevents the *Ae. aegypti* oviposition activity.

## Introduction

*Aedes aegypti*, vector of Dengue, Chikungunya and Zika viruses, is a species spread widely throughout the World and successful synanthropic mosquitoes, taking advantage of different breeding sites similar to those used in its original habitat [1]. The establishment of populations of this species depends on the invaded area characteristics, which change according to the area [2]. Tropical areas are the most affected by this vector. However, the distribution range of both the vector and pathogens transmitted has extended to temperate areas, affecting the environmental conditions not only their biology but also their distribution [3, 4].

The environment can be characterized by satellite and meteorological variables obtained from satellite images and weather stations, respectively, over time [5]. Information of satellite variables has been used in different studies to characterize the environment where the vector is settled on, to assess its influence on *Ae. aegypti* throughout the year and to develop models. In several studies, the importance of variables has been demonstrated, being the Land Surface Temperature (LST) and the Normalized Difference Vegetation Index (NDVI) both of the most used variables in *Ae. aegypti* models developed [6–8].

On the other hand, the meteorological variables are good *Ae. aegypti* abundance predictors. Temperature is one of the variables mainly used in predicted models since it influences on the life cycle of these mosquitoes, adult longevity, female fecundity [9], vectorial capacity [10], adult size, immature development, as well as for survival rates [11, 12]. Minimum temperature is related in a positive way with the vector abundance [13], what affect the flight performance [14]. In general, the precipitation is also associated positively with *Ae. aegypti* abundance [15] since it mainly fill containers what increase the number of breeding sites and directly the abundance of the vector [16]. Furthermore, there is evidence of the impact that these meteorological variables have on the increase of the diseases incidence transmitted by *Ae. aegypti* such as dengue [17, 18].

In this study, vegetation indices derived from satellite images were used, and meteorological variables, such as temperature, precipitation, humidity, vapor pressure of water and photoperiod obtained from weather stations. Based on the results found in the aforementioned studies, it was expected that temperature and precipitation affect *Ae. aegypti* oviposition activity. In that case, when temperature and precipitation increase, it would be expected an increase in the number of laid eggs weeks after. A similar relationship was expected with the vegetation indices, since temperature and precipitation regulate, to some extent, the vegetation development [19].

Despite the importance of the variables mentioned above, the vector activity fluctuates differently according to the study area and the sampling scale, mainly in areas where the vector is not present throughout the year, as well neither the occurrence of dengue cases [2, 20]. For this reason, it is important to study the vector activity in each particular area or region throughout the year in order to develop the best management strategies. *Aedes aegypti* surveillance using ovitraps is one of the most effective and low cost methods to know its distribution and the seasonal fluctuation of the populations [21]. Predictive models based on mentioned variables that characterize the environment and determine the vector dynamics are useful tools that help in the surveillance and control of these mosquitoes; also help to assess the outbreak risk at any area at a certain time. Therefore, the objective of this work is to evaluate *Ae. aegypti* oviposition activity in relation to satellite and meteorological variables in Córdoba City (Argentina), through the development of temporal generalized linear mixed models.

## Material and Methods

### Ethics statement

Ovitrap sampling was carried out under the *Ae. aegypti* surveillance program of the Córdoba Province Ministry of Health so no written informed consent was required. The residents of the dwellings where ovitraps were placed provided oral informed consent.

### Study area

The study took place in Córdoba City (31° 24’ S, 64° 11’ W) of Córdoba Province, Argentina, belonging to a temperate region (Fig 1). The cold and dry period is from May to September, while the warmest and rainy period is from October to April. According to the records from October 2010 to April 2017, the mean annual rainfall is 800 mm; during the cold period, the maximum and minimum average temperatures are 20°C and 8°C, while in summer are 29°C and 16°C, respectively [22].

**Fig 1.**
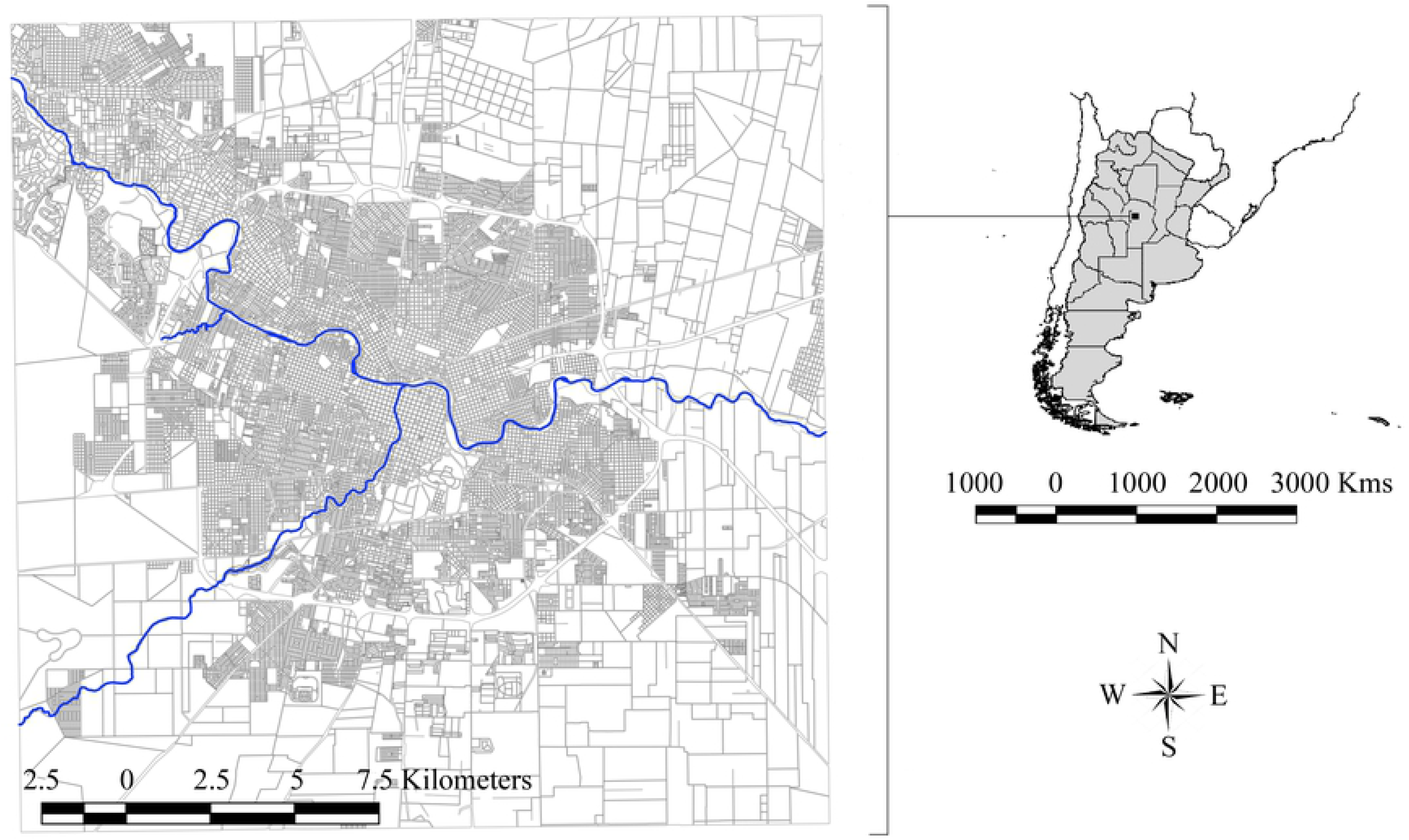
Study area where the samplings were carried out in Córdoba City, from 2009 to 2012.

### Ovitrap sampling

*Aedes aegypti* oviposition activity fluctuation was studied during three vector activity seasons, from November to May (2009-2010: Season 1, 2010-2011: Season 2, 2011-2012: Season 3), and the fluctuation was assessed weekly by ovitrap sampling. The study was carried out in 177 dwellings uniformly distributed throughout Córdoba City. An ovitrap was placed outdoors each dwelling in a shade area, protected from the rain and located at ground level. Ovitraps consisted of 350 ml transparent plastic jars (8 cm in diameter and 13 cm high), with a cylinder of brown heavy (120 g) rough paper inside that cover the entire interior wall. Ovitraps were filled with 250 ml of grass infusion when settled. This infusion consisted in dry grass macerated in water during a week, which works as an attractant to gravid *Ae. aegypti* females [23]. Ovitraps were replaced weekly. At the laboratory, laid eggs by ovitrap were counted. The response variable used was the weekly average number of eggs per active ovitraps, considering non-active traps those absent or destroyed [24]. Sampling was carried out by the Department of Epidemiology of the Córdoba Province Ministry of Health in cooperation with the mosquito team of the Instituto de Investigaciones Biológicas y Tecnológicas, IIBYT (CONICET-Universidad Nacional de Córdoba).

### Satellite and meteorological variables

Vegetation indices from MODIS products were used as satellite variables. NDVI (MOD13Q1.006), EVI (MOD13Q1.006) and LAI (MOD15A2H.006) were retrieved from the online Application for Extracting and Exploring Analysis Ready Samples (AppEEARS), courtesy of the NASA EOSDIS Land Processes Distributed Active Archive Center (LP DAAC), USGS/Earth Resources Observation and Science (EROS) Center, Sioux Falls, South Dakota (https://lpdaacsvc.cr.usgs.gov/appeears/). In Table 1, the satellite variables with their characteristics are shown (https://lpdaac.usgs.gov/dataset_discovery/modis/modis_products_table) [25].

**Table 1.** Satellite variables obtained from MODIS products.

| **Variable** | **Products** | **Spatial**<br>**resolution** | **Temporal**<br>**resolution** | **Description** |
| --- | --- | --- | --- | --- |
| **NDVI**<br>(Normalized<br>Difference<br>Vegetation<br>Index) | MOD13Q1.006 | 250 meters | 16 days | Chlorophyll sensitive<sup>[25]</sup> |
| **EVI**<br>(Enhanced<br>Vegetation<br>Index) | MOD13Q1.006 | 250 meters | 16 days | Responsive to canopy<br>structural variations (leaf<br>area index, canopy type,<br>plant physiognomy, and<br>canopy architecture)<sup>[25]</sup> |
| **LAI** (Leaf<br>Area Index) | MOD15A2H.006 | 500 meters | 8 days | Defined as the one-sided<br>green leaf area per unit<br>ground area in broadleaf<br>canopies and as one-half the<br>total needle surface area per<br>unit ground area in<br>coniferous canopies. |

Data were obtained from Córdoba City, from September 6, 2009 to May 26, 2012. There were a few gaps in the data, because of frequency of the satellite and the variables were not measured. When that occurred, the variables were extended to weekly time series doing data interpolation to merge with *Ae. aegypti* eggs count data.

Meteorological National Service (MNS) provided daily and hourly measurements, for the sampling period, from the two weather stations placed for Córdoba City by the National Institution. Weekly meteorological data (maximum/minimum temperature, temperature range, accumulated rainfall, mean relative humidity, humidity range and mean vapor pressure of water) were calculated and used to data analysis. Besides, the photoperiod was added as variable, which has an effect on lifespan and blood feeding activity and controls the species survival [26]. Considering the biological characteristics of *Ae. aegypti*, time lags were applied to the explanatory variables [7, 27], meaning the length of time that has elapsed since the change in weather conditions that is reflected in the response variable in order to obtain a better fitted model.

### Data analysis

Satellite and meteorological variables were standardized by subtracting their means and dividing by their standard deviations. Simple linear correlation analysis was performed with time lags from 1 to 6 weeks between the response variable and the explanatory variables, using Spearman’s correlation coefficient. Variables with the time lags that best correlated with the response variable were chosen to fit the models. The average number of weekly eggs per active ovitraps, of the 177 ovitraps, was modeled as a function of the set of satellite and meteorological variables, with the selected time lags from the first (November 2009-May 2010) and second (November 2010-May 2011) sampling seasons of study, using Generalized Linear Mixed Models (GLMM). Weeks were included as random effect to incorporate the temporal dependency. In the first place, decision tree and univariate models were performed to choose most important variables and start to develop the models, as well as, identifying possible interactions between variables. Five specific models were set up which were performed in Statistical software *R* v.3.3.1 (R Core Team 2016), assuming a negative binomial distribution of errors with log link function. Variable significance was evaluated and when there were not significant variables, interaction between the significant variables were added to improve the model [28]. The multicollinearity between variables was evaluated in the models. Besides, overdispersion and the normality of the distribution of residuals were checked. Models were ranked following the Akaike’s Information Criterion (AIC) and that with the lowest AIC was selected as the best model [29]. Finally, we assessed whether our selected model was a good predictor of the average number of laid eggs by *Ae. aegypti* during the third sampling season (November 2011-May 2012). The Spearman’s correlation coefficient was calculated to assess the relation between the values observed in the third sampling season and those estimated by the model in order to validate it.

## Results

### Entomological data and temporal analysis

A total of 268,874 *Ae. aegypti* eggs were collected during the three sampling seasons, recording 101,278 in the first sampling season, 65,640 in the second and 101,956 in the third sampling season. During the first sampling season, two peaks of oviposition activity occurred in mid-January and at the end of February-beginning of March (Fig 2). Moreover, maximum (38.8°C) and minimum (20.7°C) temperatures were observed during the previous fifth week to the second peak of oviposition activity. In addition, the precipitation was the largest in the previous third week to the maximum peaks of *Ae. aegypti* oviposition activity, as well as humidity showed a similar pattern. In the second sampling season, the number of eggs increased reaching a maximum in mid-January, then diminished slightly and fluctuated until mid-March, declining abruptly at the end of the sampling season. Likewise, the maximum temperature fluctuated slightly during this sampling season maintaining above 30°C without showing a noticeable peak, while the variation of minimum temperature was similar the maximum varying between 7.5°C and 19.9°C. Regarding precipitation, even though there was not important peaks before the greatest oviposition activity, there was large precipitations in the following weeks which was reflected in a slightly increase of the humidity with passing of time. During the last sampling season, a noticeable maximum of activity was observed in the middle of March with maximum temperatures above 30°C and minimum temperature between 9.3°C and 18.8°C during the previous weeks to the peak of oviposition activity. Concerning precipitation and humidity, a similar pattern to the first sampling season was observed which means previous to the maximum of oviposition large rainfalls and humidity increased.

**Fig 2.**
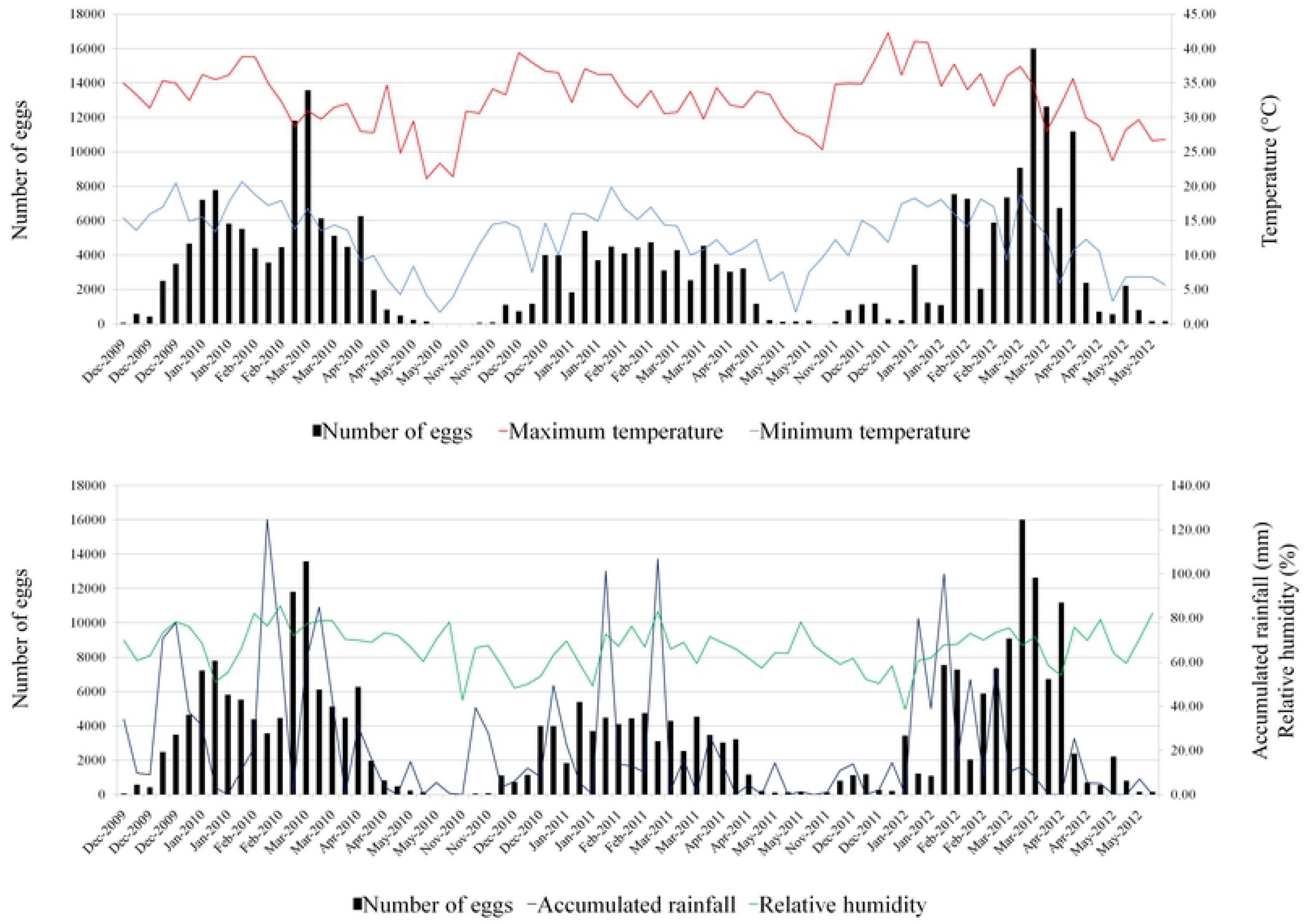
Fluctuation of egg number and meteorological variables from 2009 to 2012, in Córdoba City (Argentina).

### Development of *Aedes aegypti* oviposition model

The explanatory variables that were selected to fit the models are shown in Table 2 with their best time lag and correlation coefficient with the response variables, which were significant (p<0.05) showing effect on oviposition activity. Those explanatory variables that were correlated greater than 0.7 were not incorporated into the same model. According to the analysis carried out, the most relevant variables were: the EVI vegetation index, accumulated precipitation, vapor pressure and minimum temperature.

**Table 2.** Variables used to develop the GLMM and time lag of the satellite and meteorological variables that best correlated with the response variable and their respective correlation coefficients.

|  | Variables | Time lags with best correlation (weeks) | Period from which the variable was obtained (days) | Spearman Correlation coefficients |
| --- | --- | --- | --- | --- |
| **NDVI<sub>16</sub>** | NDVI | 4 | 16 | -0.395 |
| **EVI<sub>16</sub>** | EVI | 3 | 16 | 0.733 |
| **LAI<sub>8</sub>** | LAI | 1 | 8 | 0.746 |
| **Tmin<sub>1</sub>** | Minimum temperature | 1 | 7 | 0.601 |
| **Tmin<sub>3</sub>** | Minimum temperature | 3 | 7 | 0.717 |
| **Tmin<sub>49</sub>** | Minimum temperature | 1 | 49 | 0.835 |
| **Tmax<sub>1</sub>** | Maximum temperature | 1 | 7 | 0.264 |
| **Trange<sub>28</sub>** | Temperature range | 1 | 28 | -0.643 |
| **RAIN<sub>28</sub>** | Accumulated rainfall | 1 | 28 | 0.767 |
| **RHmin<sub>28</sub>** | Minimum relative humidity | 1 | 28 | 0.481 |
| **RHmean<sub>21</sub>** | Mean relative humidity | 1 | 21 | 0.45 |
| **RHrange<sub>35</sub>** | Relative humidity range | 1 | 35 | -0.513 |
| **VP<sub>28</sub>** | Mean vapor pressure of water | 1 | 28 | 0.886 |
| **Ph<sub>28</sub>** | Photoperiod | 1 | 28 | 0.52 |

The obtained models are shown in Table 3. The explanatory variables that were included in the models were significantly important and multicollinearity was not found. The best model M2 (AIC= 332.5) included EVI vegetation index, vapor pressure of water, precipitation and photoperiod. Based on this model, a positive relationship is observed between EVI vegetation index, vapor pressure of water, photoperiod and the response variable while the interaction between vapor pressure of water and photoperiod was negatively associated with the number of eggs, as well as precipitation. According to the model obtained, for each increment unit in EVI vegetation index, the average number of eggs increases by 72% and for each increase in vapor pressure (hPa), the number of eggs increases by 20%. In addition, the average number of eggs decreases by 50% for each millimeter of rain and increases by 29% for each light hour that is added.

**Table 3.** Additional information about the developed models ranked by their AIC scores.

| Model | Variables | Estimates | p-value | 95% confidence interval | Standard error | AIC |
| --- | --- | --- | --- | --- | --- | --- |
| **M1** | Intercept | 2.23 | < 2e-16 | 1.788 ; 2.632 | 0.21 | 444.9 |
|  | Weeks |  |  | 1.131 ; 1.854 |  |  |
| **M2** | Intercept | 2.211 | < 2e-16 | 2.018 ; 2.383 | 0.092 | 332.5 |
|  | EVI<sub>16</sub> | 0.54 | 3.96e-06 | 0.317 ; 0.787 | 0.117 |  |
|  | VP<sub>28</sub> | 1.201 | < 2e-16 | 0.933 ; 1.495 | 0.141 |  |
|  | RAIN<sub>28</sub> | -0.404 | 0.001 | -0.663 ; -0.166 | 0.124 |  |
|  | Ph<sub>28</sub> | 0.256 | 0.008 | 0.06 ; 0.447 | 0.097 |  |
|  | VP<sub>28</sub>*Ph<sub>28</sub> | -0.552 | 1.15e-05 | -0.813 ; -0.313 | 0.126 |  |
|  | Weeks |  |  | 0.221 ; 0.457 |  |  |
| **M3** | Intercept | 2.329 | < 2e-16 | 2.141 ; 2.495 | 0.088 | 335.6 |
|  | EVI<sub>16</sub> | 0.962 | 4.92e-11 | 0.684 ; 1.27 | 0.146 |  |
|  | VP<sub>28</sub> | 1.231 | < 2e-16 | 1.019 ; 1.471 | 0.112 |  |
|  | Trange<sub>28</sub> | 0.382 | 7.41e-05 | 0.188 ; 0.578 | 0.096 |  |
|  | EVI<sub>16</sub>*VP<sub>28</sub> | -0.7 | 1.34e-06 | -1.007 ; -0.426 | 0.145 |  |
|  | Weeks |  |  | 0.213 ; 0.467 |  |  |
| **M4** | Intercept | 2.147 | < 2e-16 | 1.927 ; 2.335 | 0.103 | 349.2 |
|  | EVI<sub>16</sub> | 0.436 | 0.0005 | 0.196 ; 0.698 | 0.124 |  |
|  | VP<sub>28</sub> | 1.198 | < 2e-16 | 0.958 ; 1.468 | 0.127 |  |
|  | RHmean<sub>21</sub> | -0.418 | 3.92e-05 | -0.629 ; -0.22 | 0.102 |  |
|  | Weeks |  |  | 0.297 ; 0.578 |  |  |
| **M5** | Intercept | 2.053 | < 2e-16 | 1.805 ; 2.266 | 0.116 | 358.3 |
|  | EVI<sub>16</sub> | 0.677 | 4.29e-06 | 0.392 ; 0.985 | 0.147 |  |
|  | Tmin<sub>49</sub> | 1.29 | 4.60e-15 | 0.978 ; 1.638 | 0.165 |  |
|  | Trange<sub>28</sub> | 0.425 | 0.002 | 0.148 ; 0.71 | 0.14 |  |
|  | Weeks |  |  | 0.378 ; 0.684 |  |  |

On the other hand, other developed models incorporated, in addition to the EVI vegetation index and the vapor pressure, variables such as temperature range, relative humidity and minimum temperature. According to these models, the average number of eggs increases by 47% and 29% for each increase in the temperature range and the minimum temperature respectively, and decreases by 52% when the relative humidity increases one unit.

Data from third sampling season was used to validate the obtained model observing a correlation coefficient of 0.84 (p<0.05) between the *Ae. aegypti* oviposition activity observed and the activity predicted by the model M2 (Fig 3).

**Fig 3.**
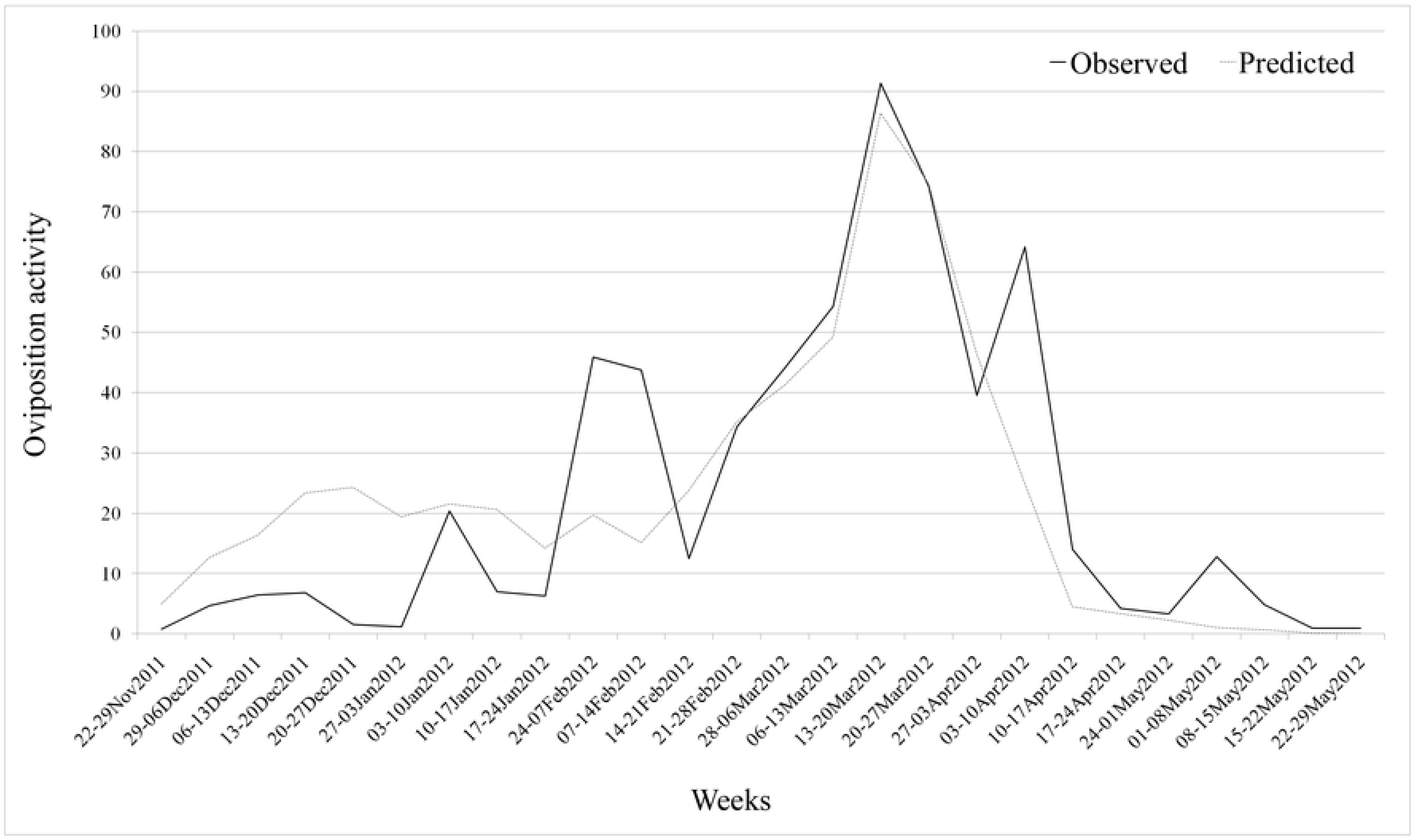
Oviposition activity observed and predicted in the third sampling season, in Córdoba City (Argentina).

## Discussion

According to the results here obtained, it is important to highlight the number of eggs was 35% lower in the second sampling season in relation to the other two seasons. The average maximum and minimum temperature were similar in the three sampling seasons, 32.6°C and 12.5 °C, respectively. Precipitation was greater in the first sampling season (758.5mm) while in the other two it was less than 500mm; however, in November of the first and third sampling seasons there were higher rain average per week (34mm and 29mm, respectively) than in November of the second (17mm) what indicates, somehow, the relevance of rain distribution during the vector activity season. The rainfall amount in each sampling season would affect the availability of containers [30], and the number of laid eggs. It is assume a great quantity of eggs that were laid in the first rainy sampling season did not be covered by water in the next scarce rainy season (second sampling season) and therefore eggs did not hatch; as a consequence, the number of adults was lower in the environment what explain the low number of eggs detected during this second sampling season. Furthermore, in this sampling season, females could have laid the eggs at a lower level in the recipients, that contained less water also related to the lower precipitations recorded, which allowed eggs to hatch in the third sampling season in spite of the scarce rainfall. In addition, the females could have chosen the ovitraps as potentially better breeding sites and, therefore, may have laid most eggs in them instead of in other artificial containers with less amount of water, what could explain the higher number of eggs found in the third sampling season.

The significant importance of satellite and meteorological variables on the oviposition activity of *Ae. aegypti* is demonstrated here. Vegetation index is one of the most important variables, showing a positive relationship and fluctuating in a similar way as the oviposition activity. This relationship has also been observed between the NDVI and the egg number fluctuation of this species in the north of the country [7]. In addition, studies in La Habana, Cuba, showed a correlation between the number of larva infested containers and the presence of trees and vegetation, which demonstrates the importance of the shade that vegetation provides to containers in which the *Ae. aegypti* develops, always depending on the container use and location [31].

At the sub-tropical climatic conditions of northern Argentina, a positive relationship was observed between *Ae. aegypti* egg fluctuation and NDVI which is directly related to precipitation from previous weeks [7]. In Córdoba City with a temperate climate and cold winters without vector activity, larval development and then the presence of adults during the favorable season depends mainly on the previous vector activity season, the laid eggs in containers and precipitations to induce hatching. The negative relationship between the average number of eggs and rainfall here obtained could be explain since containers filled with water, due to the abundant rains, avoid females to lay eggs. In Malaysia was observed that *Ae. albopictus* females preferred to lay eggs in half-filled containers instead of in overflow containers because these acts as a repellent for gravid females. This is because the rains tend to fill the containers and spill the water that is in them, and consequently this eliminates the larvae that are developing in those breeding sites [32]. Another possible explanation for this negative relationship may be that when the rains are not abundant, the human population tends to store water which means that there is availability of breeding sites for the vector although rains are scarce [33]. Maybe other explanation is applicable for Córdoba City, but it is remarkable the importance of precipitation peaks that usually occur weeks before finding eggs, adult mosquitoes or dengue cases appear in coincidence with the results of other researchers [27, 34, 35].

In temperate zones, the minimum temperature can act as a limiting factor, slowing oviposition activity as happens in Buenos Aires [34], and Córdoba [36] not only during the winter season, resulting of greater importance than the maximum temperature in this study. Both the minimum temperature and precipitations were of the most important variables in this study, in coincidence with other studies [27, 37], however, these two variables were not incorporated into a model because they are highly correlated. In relation to maximum temperature, a positive relationship is observed with the oviposition activity. This relationship is not linear because very high temperatures can affect the survival of the vector in the case of the tropics for instance [35], what does not occurs in Córdoba since maximum temperatures are not usually extremely high, reason by which could be applicable a linear relationship for our study area. This would explain in some way why the maximum temperature, despite having a significant correlation, did not enter into any of the developed models. It is also important to consider the thermal amplitude since the temperature fluctuation has a negative effect on *Ae. aegypti* in Thailand [38], observing fewer eggs when the thermal amplitude is greater (November and May in the case of Córdoba), as was observed in the correlations. However, in the models this variable turned out to show a positive relationship with the response variable maybe because of the interaction that has with other variables.

Estallo et al. [27] found a positive relationship between *Ae. aegypti* egg number and the vapor pressure of water showing a decrease in eggs when this variable decreases its value. Similar to this, a high correlation was found here between these two variables with a positive association entering this variable in most of our models. This could be closely related to the humidity which is associated to the vapor pressure of water. Additionally, there are works that point out the importance of the relative humidity on the vector activity despite its low correlation, and when all conditions are given the activity of the species is low since the relative humidity is not adequate [27, 30]. In our work, although this variable showed a low correlation value, it was important since it entered in one of our models.

The photoperiod was not one of the main variables to develop the models, however, it was important in association with variables like EVI vegetation index, vapor pressure of water and precipitation, contradicting that observed in Orán (northern Argentina), where the photoperiod had not a significant effect on the vector [27]. This contradiction may be due to the difference that exists in terms of climatic conditions indicating the difference in the effect of the same variables in different latitude, evidencing spatial variations and that for this reason care must be taken when results are extrapolated from one place to another [35].

## Conclusions

Satellite and meteorological variables are key factors explaining the variability of egg number of *Ae. aegypti*, resulting in this study the predominant variables for our temperate region those shown in the model M2: EVI vegetation index, vapor pressure of water, precipitation and photoperiod, what was partially expected since temperature was not included in the best model, although vegetation index is directly related with both, temperature and precipitation. Water vapor pressure and photoperiod are also variables that should be more consider in this kind of studies. Our work will help to clarify the activity of *Ae. aegypti* in temperate regions, since the results are not the same compared with those of tropical regions, highlighting the importance of local studies and moreover in temperate areas such as Córdoba, and other temperate areas of the globe where not much studies have focused.

## Acknowledgments

The authors thank the personnel of Epidemiological area of the Province Health Ministry of Córdoba for their support on the field sampling and collaboration obtaining data for this study. ELE and WRA are members of the researcher career of National Scientific and Technical Research Council (CONICET) - Argentina, and EMB has a doctoral scholarship from CONICET as a PhD student, whose thesis is under the direction of ELE and FLA. We thank the Consejo Nacional de Investigaciones Científicas y Tecnológicas (CONICET) and the Proyecto de Investigación de Unidades Ejecutoras CONICET 2016 (entitled “Evaluación eco-epidemiológica de vectores de arbovirus en ecosistemas urbano-periurbanos a través de un enfoque transdisciplinar para el desarrollo de estrategias de manejo integrado”) of which EMB is a fellow.

